# Sputnik virophage disrupts the transcriptional regulation of its host giant virus

**DOI:** 10.1101/2024.08.09.607287

**Authors:** Jingjie Chen, Hiroyuki Ogata, Hiroyuki Hikida

## Abstract

Sputnik virophages are small double-stranded DNA (dsDNA) viruses that replicate only inside host amoebae infected with giant dsDNA viruses, mimiviruses. Sputnik infection affects mimivirus replication, but their molecular interaction remains poorly understood. Here, we performed a time-course transcriptome analysis of *Acanthamoeba castellanii* cells infected with Acanthamoeba polyphaga mimivirus (APMV) (hereafter referred to as Sputnik^−^ cells) and those infected with both APMV and Sputnik 3 virophage (Sputnik^+^ cells). The gene expression patterns of the amoebae were similar between these two conditions, whereas the expression of APMV genes was drastically affected by Sputnik, depending on the timing of their expression. Early expressed APMV genes showed similar expression patterns in both conditions at the early stage of infection. However, at late stages, their expression levels remained higher in Sputnik^+^ cells than in Sputnik^−^ cells, suggesting a prolongation of early gene expression by Sputnik. Late expressed APMV genes showed lower expression at earlier stages in Sputnik^+^ cells, but their expression levels reached or exceeded those in Sputnik^−^ cells at later stages, indicating a delay of gene expression. Overall, our results demonstrated that Sputnik infection drastically alters the transcriptome of APMV rather than amoeba, likely by disturbing the transition from early to late stages of APMV infection.

## Introduction

Virophages are double-stranded DNA (dsDNA) viruses associated with other dsDNA viruses, nucleocytoviruses, which have large genomes and particles, and are often called as giant viruses^1–3^. The first virophage, Sputnik, has an 18 kbp genome in 80 nm particles and is associated with a giant virus, mimivirus, which has a 1.2 Mbp genome in 700 nm particles. Sputnik and mimiviruses infect free-living amoebae of the genus *Acanthamoeba*, but Sputnik cannot replicate without mimivirus infection. Furthermore, Sputnik infection reduces the infectivity of progeny mimiviruses, leading to the concept of “a virus infecting another virus” and the designation of virophage^4^.

Following the discovery of Sputnik, related viruses have been isolated with mimiviruses and other nucleocytoviruses^1,5^, such as Mavirus associated with Cafeteria roenbergensis virus (CroV)^6^ and Gezel-14T associated with Phaeocystis globosa virus-14T (PgV-14T)^7^. In addition to the virophages isolated with these giant viruses, sequences of virophage relatives have been identified in various environmental metagenomic data,^8–10^ ranging from aquatic environments^11^ to the human gut rumen^12^.

Not only Sputnik, but multiple isolated virophages also affect host nucleocytovirus replication by reducing progeny production^7,13–17^. However, the interaction between virophages and host nucleocytoviruses varies depending on the combinations. Sputnik virophages are thought to attach to the surface fibrils of host mimiviruses and be incorporated into host amoeba cells together^18^. In contrast, Mavirus, a virophage of CroV, can enter host protist cells independently and wait for the corresponding giant virus infection^6,17^. Among the virophages infecting mimiviruses, Zamilon virophages replicate without affecting host virus replication^16^, unlike Sputnik virophages.

The expanded number of isolates and genomic sequences discovered from metagenomic data has provided novel insights into the vast diversity of this complex tripartite system. However, the molecular mechanisms underlying these microbial eukaryote-giant virus-virophage interactions remain largely unknown. In the present study, we focused on one of the tripartite systems, *A. castellanii*, Acanthamoeba polyphaga mimivirus (APMV), and Sputnik 3 virophage, analyzing their transcriptome dynamics. Our results strengthen the concept of virophages as viruses infecting other viruses and identify the disturbance of APMV transcriptional regulation caused by Sputnik infection.

## Results

### Transcriptome landscape of the tripartite system

Since the infection cycle of Sputnik was not fully explored, we first determined the time points for RNA sequencing based on the expression timing of marker genes: the DNA polymerase B (*polB*) gene of APMV and the major capsid protein (*mcp*) genes of both APMV and Sputnik (Supplementary Fig. 1). The expression of APMV *polB* peaked at 3 hours post infection (hpi), while the expression of APMV *mcp* peaked at 6 hpi and was maintained until 9 hpi. Sputnik *mcp* showed a similar pattern to APMV *mcp*. At 12 hpi, *polB* expression reactivated and *mcp* expression decreased, suggesting that the first infection cycle ends, and secondary infection starts at this point. Based on these results, we performed transcriptome analysis at 0, 3, 6, and 9 hpi using amoeba cells infected only with APMV (Sputnik^−^ cells) and those infected with both APMV and Sputnik (Sputnik^+^ cells).

In both conditions, the proportion of amoeba-derived reads decreased while the reads from viruses increased throughout the infection (Fig. 1). In Sputnik^−^ cells, the proportion of APMV-derived reads reached approximately 50% at 9 hpi, whereas in Sputnik^+^ cells, the APMV and Sputnik reads accounted for about 20% at 9hpi. The total proportion of viral reads in Sputnik^+^ cells was around 40%, similar to the viral-read proportion in Sputnik^−^ cells.

**Fig 1.**
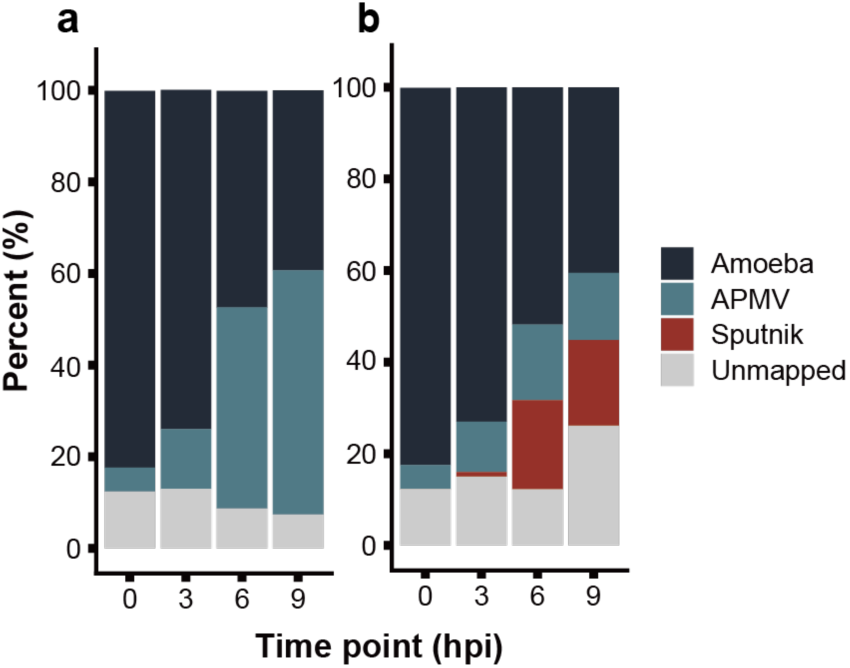
Proportions of viral and host reads at different time points. Average proportion of reads from different sources relative to total reads (a) Sputnik^−^ cells, where amoeba cells were infected only with APMV. (b) Sputnik^+^ cells, where amoeba cells were infected with both APMV and Sputnik. (n = 3).

### Sputnik genes are divided into two groups by their expression timing

As Sputnik gene expression was very low at 0 hpi (Supplementary Table 1), we investigated its gene expression at 3, 6, and 9 hpi. Sputnik genes clustered into two groups based on their expression patterns (Fig. 2). The first group, early genes (5 genes), exhibited high expression at 3 hpi, which gradually decreased from 6 to 9 hpi. The second group, late genes (17 genes), had low expression at 3 hpi but increased at 6 hpi and remained at a similar level until 9 hpi (Fig. 2a). Early genes included a gene annotated with a DNA replication function, while late genes included virion-related genes such as a DNA packaging protein, a membrane protein, and capsid proteins (Supplementary Table 1).

**Fig 2.**
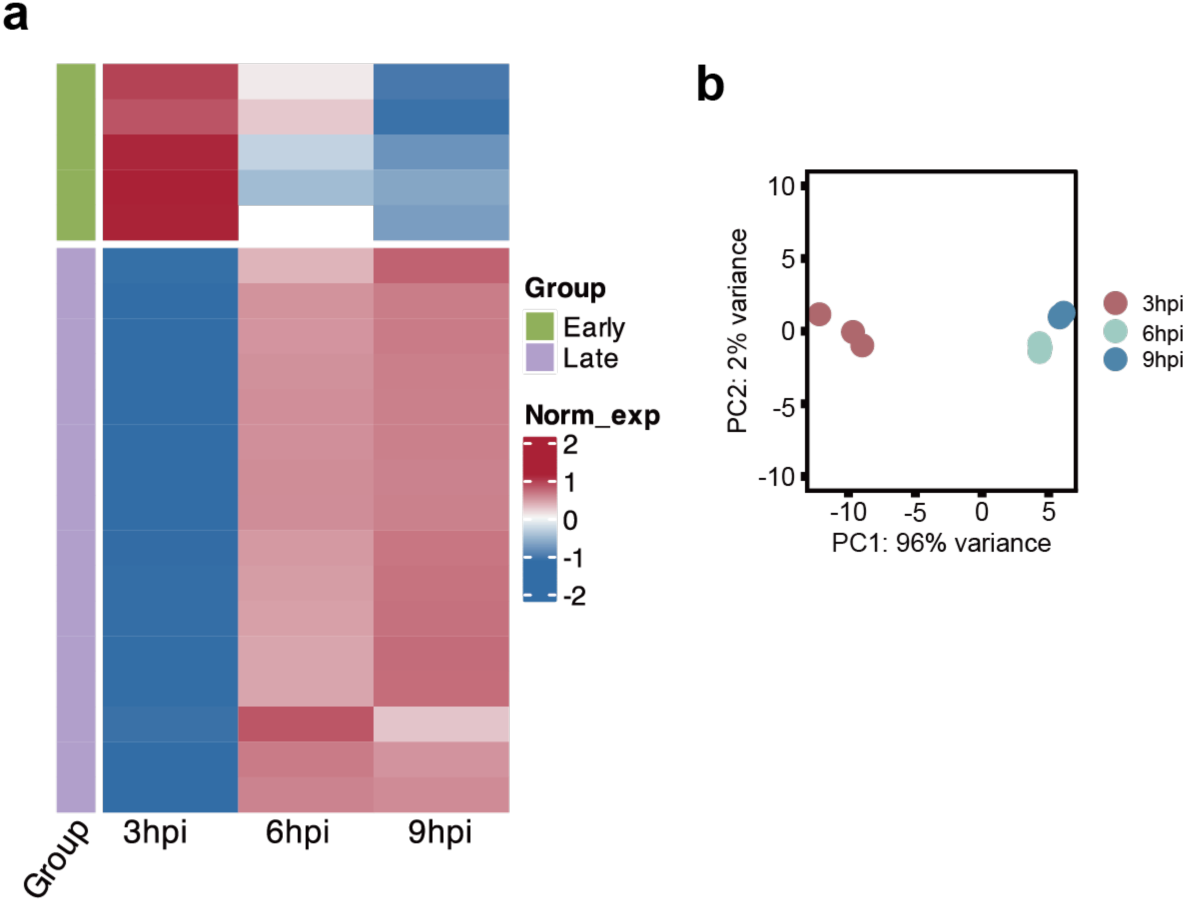
Gene expression pattern of Sputnik. (a) Heatmap and clustering based on normalized Sputnik gene expression. Gene expression was calculated as transcripts per million (TPM) and further normalized within each gene. In the left color bar, green and purple indicate early and late genes, respectively. (b) Principal component analysis (PCA) of Sputnik gene expression. Each circle represents a replicate, and each color represents a time point (n = 3).

### Sputnik affects a late stage of APMV infection

Principal component analysis (PCA) based on APMV gene expression showed that samples from Sputnik^−^ and Sputnik^+^ cells were closely located at 0 and 3 hpi (Fig. 3a). At these two time points, zero and 130 APMV genes were differentially expressed, respectively (Fig. 3b). These results suggest similar transcription profiles of APMV at the early stages of infection between Sputnik^−^ and Sputnik^+^ cells. However, at 6 and 9 hpi, samples from Sputnik^−^ and Sputnik^+^ cells were distinctly separated (Fig. 3a). Of the 979 APMV genes analyzed, 412 (42%) and 402 (41%) genes were differentially expressed at these later time points, respectively (Fig. 3b). Samples from Sputnik^−^ cells at 6 hpi overlapped with those from Sputnik^+^ cells at 9 hpi in the PCA (Fig. 3a), suggesting that Sputnik infection inhibits the progression of APMV gene expression.

**Fig 3.**
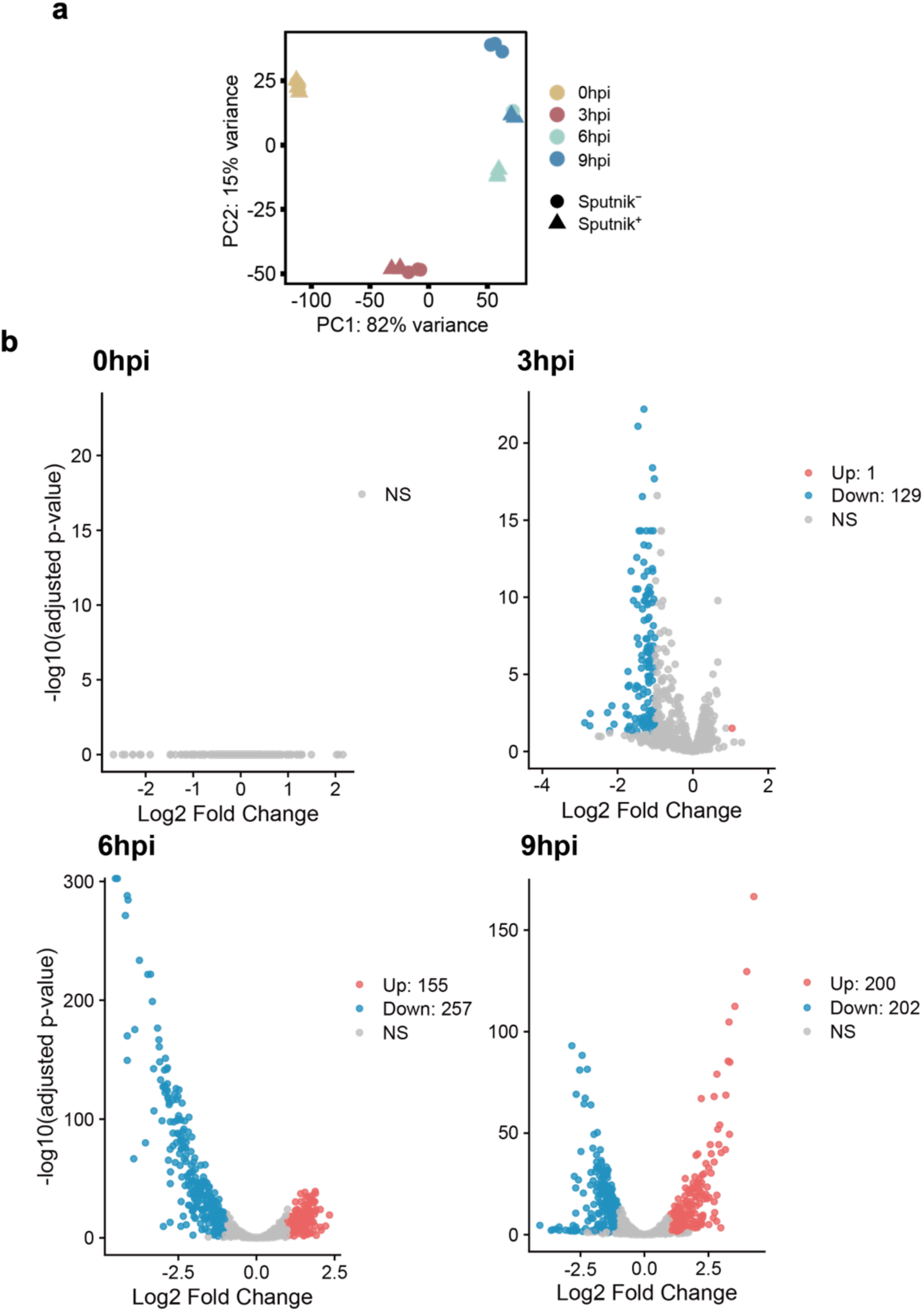
Comparison of APMV gene expression level between Sputnik^−^ and Sputnik^+^ cells. (a) PCA of APMV gene expression. Colors represent each time point, with circles and triangles representing replicates from Sputnik^−^ cells and Sputnik^+^ cells, respectively. (b) Expression changes of APMV genes are shown in volcano plot for each time point. Sampling times are indicated at the top left. Genes with an adjusted *p-* value (*p_adj_*) ≤ 0.05 and |log2 fold change| ≥ 1 are considered as differential expression genes (DEGs) and are marked in red or blue. Red and blue indicates up-regulated and down-regulated genes, respectively. NS: no significant difference.

### Sputnik infection has little effect on amoeba gene expression

In PCA based on amoeba gene expression, samples from Sputnik^−^ and Sputnik^+^ cells at each time point were closely located (Fig. 4a). No amoeba genes were differentially expressed at 0 and 3 hpi, while 90 (0.6%) and 98 (0.7%) amoeba genes were differentially expressed at 6 and 9 hpi, respectively (Fig. 4b and Supplementary Table 2). Compared with the number of amoeba genes whose expression levels were altered by APMV infection (Supplementary Fig. 2), these numbers were relatively small. Thus, we concluded that Sputnik infection has little effect on amoeba gene expression.

**Fig 4.**
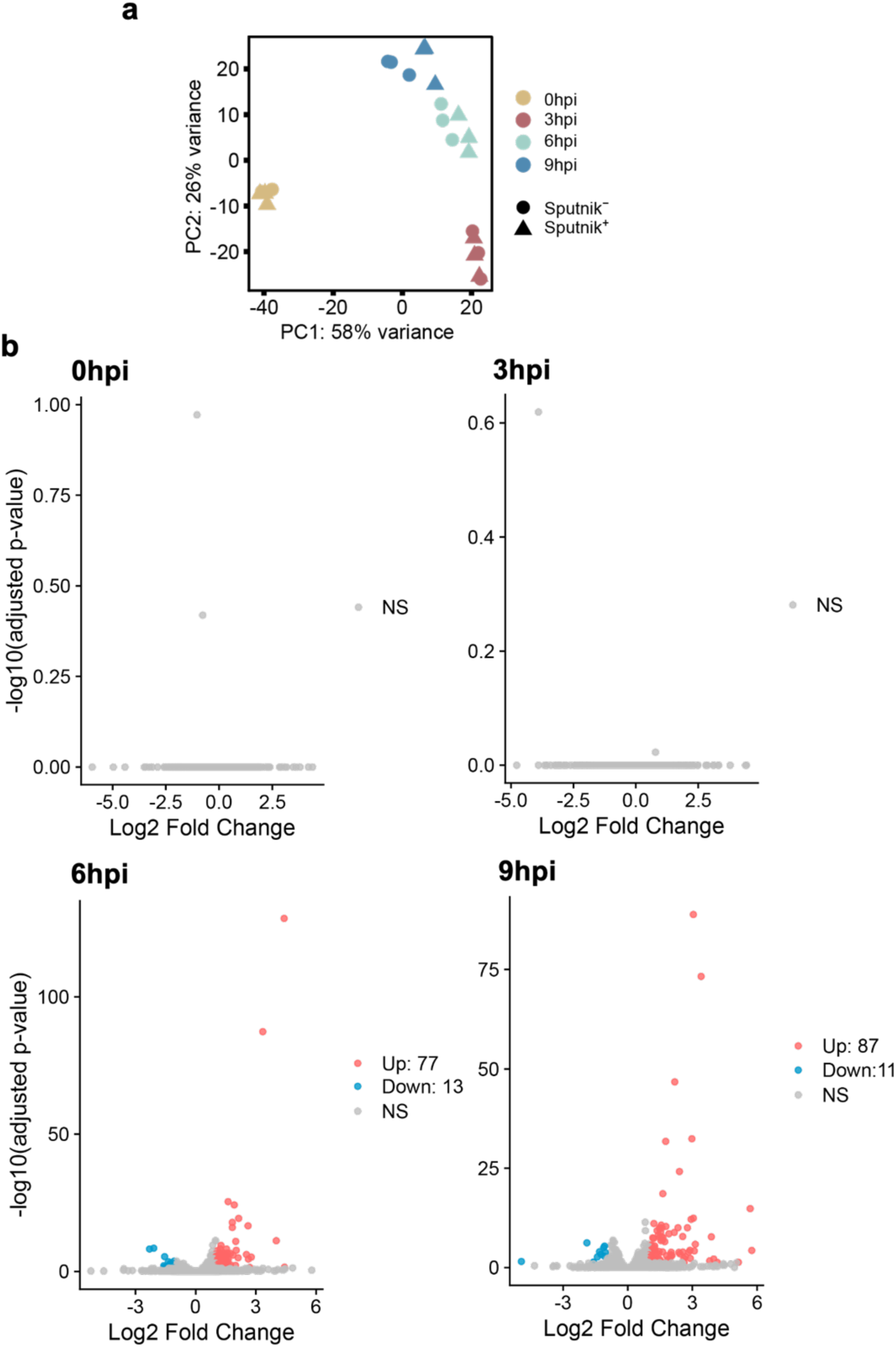
Comparison of amoeba gene expression levels between Sputnik^−^ and Sputnik^+^ cells. (a) PCA of amoeba gene expression. Circles and triangles represent each replicate from Sputnik^−^ and Sputnik^+^ cells, respectively. (b) Expression changes of amoeba genes shown by volcano plot. Amoeba gene expression was compared between Sputnik^−^ cells (as the baseline) and Sputnik^+^ cells at each time point. Genes with *p_adj_* ≤ 0.05 and |log2 fold change| ≥ 1 are defined as DEGs and are marked in red or blue. Red and blue indicates up-regulated and down-regulated genes, respectively. NS: no significant difference.

### Sputnik delays the expression of APMV genes

To elucidate the effect of Sputnik infection on APMV gene expression, we classified APMV genes based on their expression patterns in Sputnik^−^ cells. To minimize noise from genes showing minor expression changes, we focused on DEGs for downstream analysis (see Materials and Methods). We identified four clusters: immediate-early, early, intermediate, and late (Fig. 5a and Supplementary Table 3). These clusters mostly correspond to previous classifications^19^. Immediate-early genes showed high expression at 0 hpi, with a gradual decline thereafter. Early genes peaked at 3 hpi, then decreased at subsequent time points; these genes mainly include those involved in host-virus interactions and DNA replication (Supplementary Fig. 3 and Supplementary Table 3). Intermediate genes increased in expression from 0 to 3 hpi, maintained high levels at 6 hpi, and decreased at 9 hpi. Late gene expression gradually increased from 6 to 9 hpi and these genes are mainly involved in virion structure and morphogenesis (Supplementary Fig. 3 and Supplementary Table 3).

**Fig 5.**
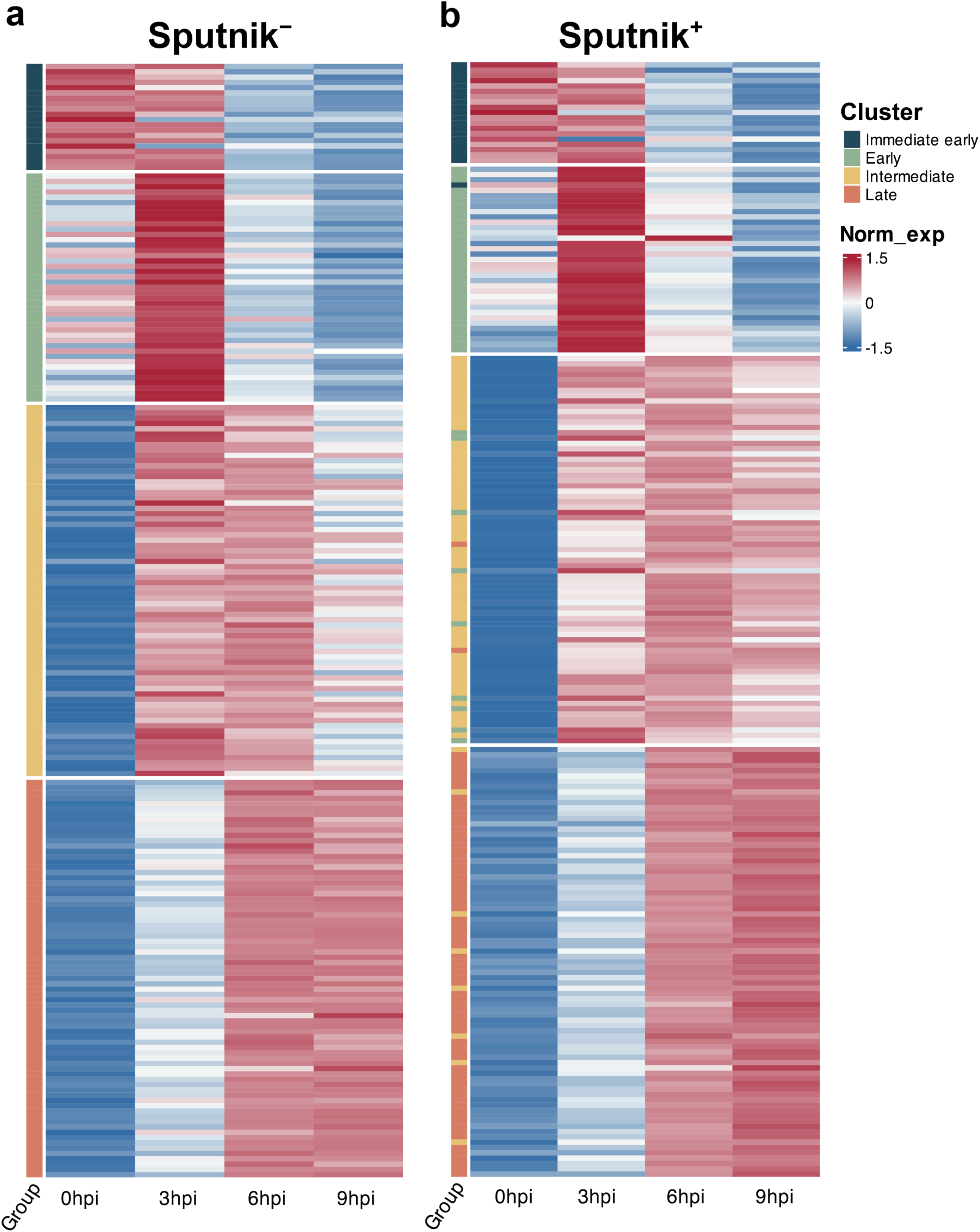
Clustering of APMV DEGs. Clustered heatmap of selected APMV DEGs from (a) Sputnik^−^ and (b) Sputnik^+^ cells. Gene expression was calculated as TPM and further normalized within each time point. Rows and columns represent genes and time points, respectively. (a) The color bar indicates each cluster. (b) The color bar indicates the clusters identified in (a).

We applied the same clustering method to the samples from Sputnik^+^ cells, resulting in four analogous clusters. While most genes clustered into the analogous clusters, some genes were grouped into clusters corresponding to later stages of expression (Fig. 5b and Supplementary Table 3). This result aligns with the delay in APMV gene expression observed in the PCA (Fig. 3a).

### Prolonged gene expression in immediate-early and early genes

In APMV genes clustered into analogous groups, some genes exhibited noticeably different expression patterns between Sputnik^−^ and Sputnik^+^ cells. For example, many intermediate genes showed relatively higher expression at 9 hpi in Sputnik^+^ cells compared to Sputnik^−^ cells (Fig. 5). To further examine APMV genes within each cluster, we performed subclustering based on the combined expression profiles in Sputnik^−^ and Sputnik^+^ cells.

In the immediate-early genes, we identified four subgroups. However, subgroup 2 and 4 comprised only two genes and one gene, respectively (Fig. 6a). Therefore, we focused on subgroups 1 and 3. Genes in both subgroups exhibited comparable expression levels in Sputnik^−^ and Sputnik^+^ cells at 0 and 3 hpi. However, genes in subgroup 1 showed approximately fourfold higher expression in Sputnik^+^ cells compared to Sputnik^−^ cells at 6 hpi, and this increased expression was maintained at 9 hpi (Fig. 6b and c). Genes in subgroup 3 also showed similar up-regulation at 6 hpi, but their expression slightly decreased at 9 hpi (Fig. 6b and c). These results suggest that Sputnik infection leads to prolonged expression of APMV immediate-early genes.

**Fig 6.**
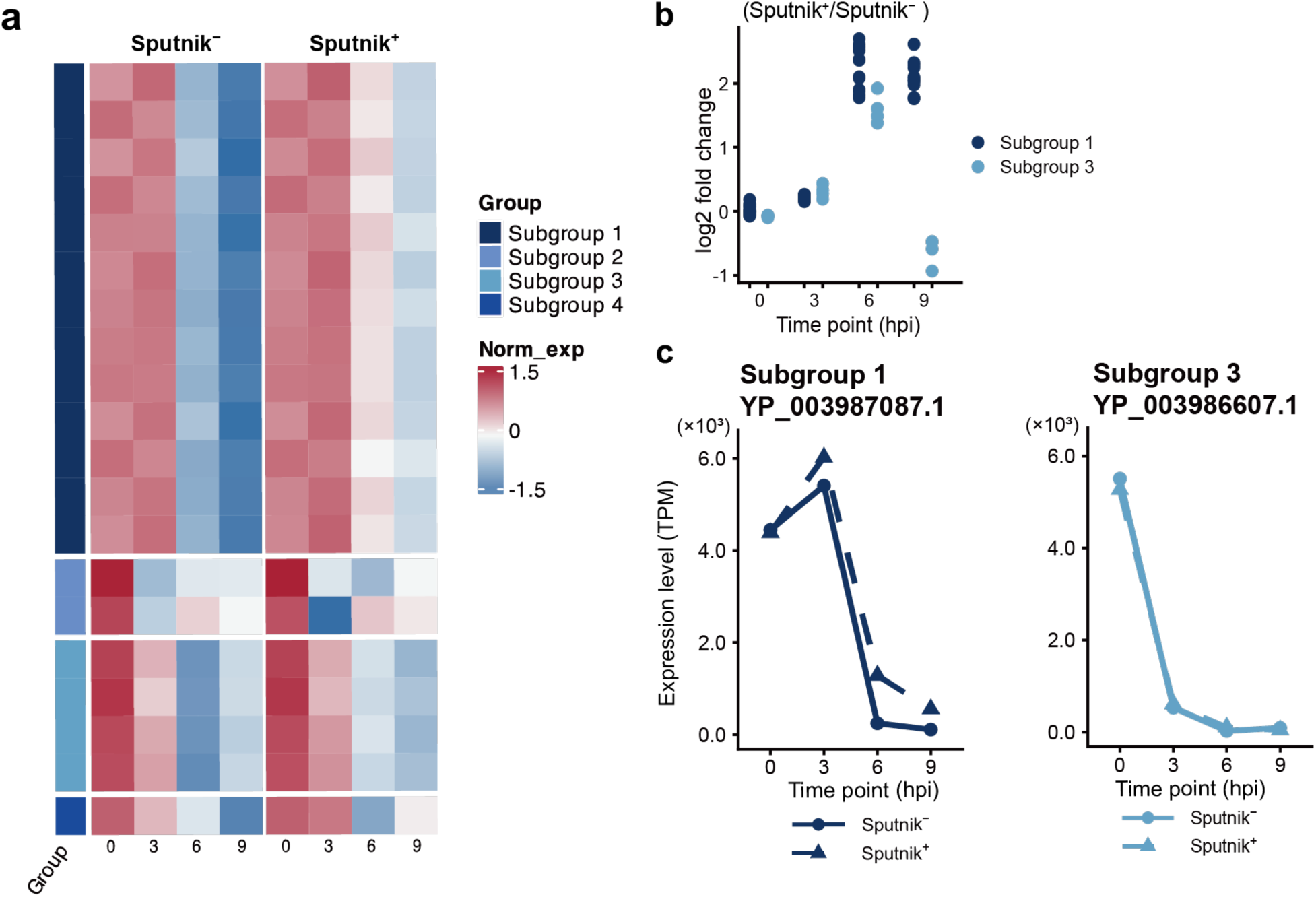
Expression profile of APMV immediate-early genes. (a) Clustered heatmap showing normalized gene expression of immediate-early genes in Sputnik^−^ and Sputnik^+^ cells. Gene expression in both Sputnik^−^ and Sputnik^+^ cells was calculated as TPM and further normalized within each gene, based on which genes were clustered. The color bar represents the subgroups of genes. (b) Expression changes from Sputnik^−^ to Sputnik^+^ cells at each time point. The log2-fold change of each gene expression was calculated based on TPM. Each dot represents an APMV gene. (c) Temporal expression changes of representative genes. Genes with the highest expression in subgroups 1 and 3 are shown.

Early genes were divided into two subgroups (Fig. 7a). Genes in both subgroups showed similar temporal expression changes. Their expression increased from 0 to 3 hpi, with similar levels in both Sputnik^−^ and Sputnik^+^ cells. However, at 6 and 9 hpi, these genes exhibited 4–8-fold higher expression in Sputnik^+^ cells compared to Sputnik– cells (Fig. 7b and c). These results suggest that Sputnik infection also prolongs the expression of APMV early genes similar to the effect observed with immediate-early genes.

**Fig 7.**
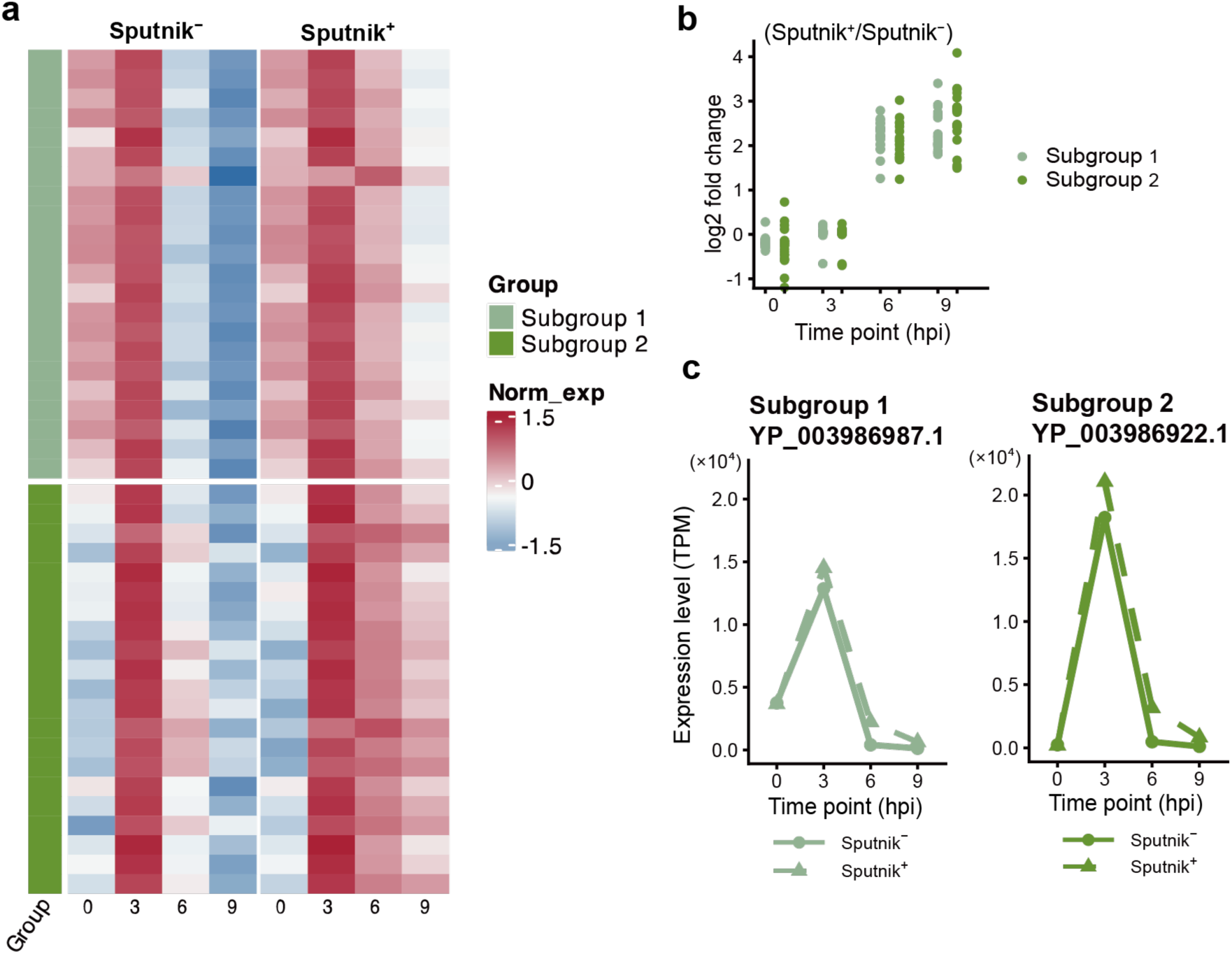
Expression profile of APMV early genes. (a) Clustered heatmap showing normalized gene expression of APMV early genes in Sputnik^−^ and Sputnik^+^ cells. Gene expression was calculated as TPM, then normalized and clustered as described in Fig.6 (b) Expression changes from Sputnik^−^ to Sputnik^+^ cells at each time point. Each dot represents an APMV gene. (c) Temporal expression changes of representative genes. Genes with the highest expression from each subgroup are shown.

### Delayed gene expression in intermediate and late genes

Intermediate genes were clustered into three subgroups (Fig. 8a). Sputnik infection resulted in slightly lower expression of genes in all three subgroups at 3 hpi, but their expression reached similar or higher level than in Sputnik^−^ cells at later stages of infection, suggesting a slight delay in the initiation of gene expression (Fig. 8b and c). Genes in subgroup 1 showed similar expression levels in both Sputnik^−^ and Sputnik^+^ cells at 6 and 9 hpi. In contrast, genes in subgroups 2 and 3 exhibited approximately fourfold higher expression in Sputnik^+^ cells compared to Sputnik^−^ cells at 6 hpi (Fig. 8b). While the expression of genes in these two subgroups decreased in Sputnik^−^ cells at 9 hpi, they maintained high expression levels in Sputnik^+^ cells, increasing the gap in expression differences between the two conditions (Fig. 8b and c). These results suggest that intermediate genes in subgroups 1 and 2 also exhibit prolonged gene expression at late stages of infection.

**Fig 8.**
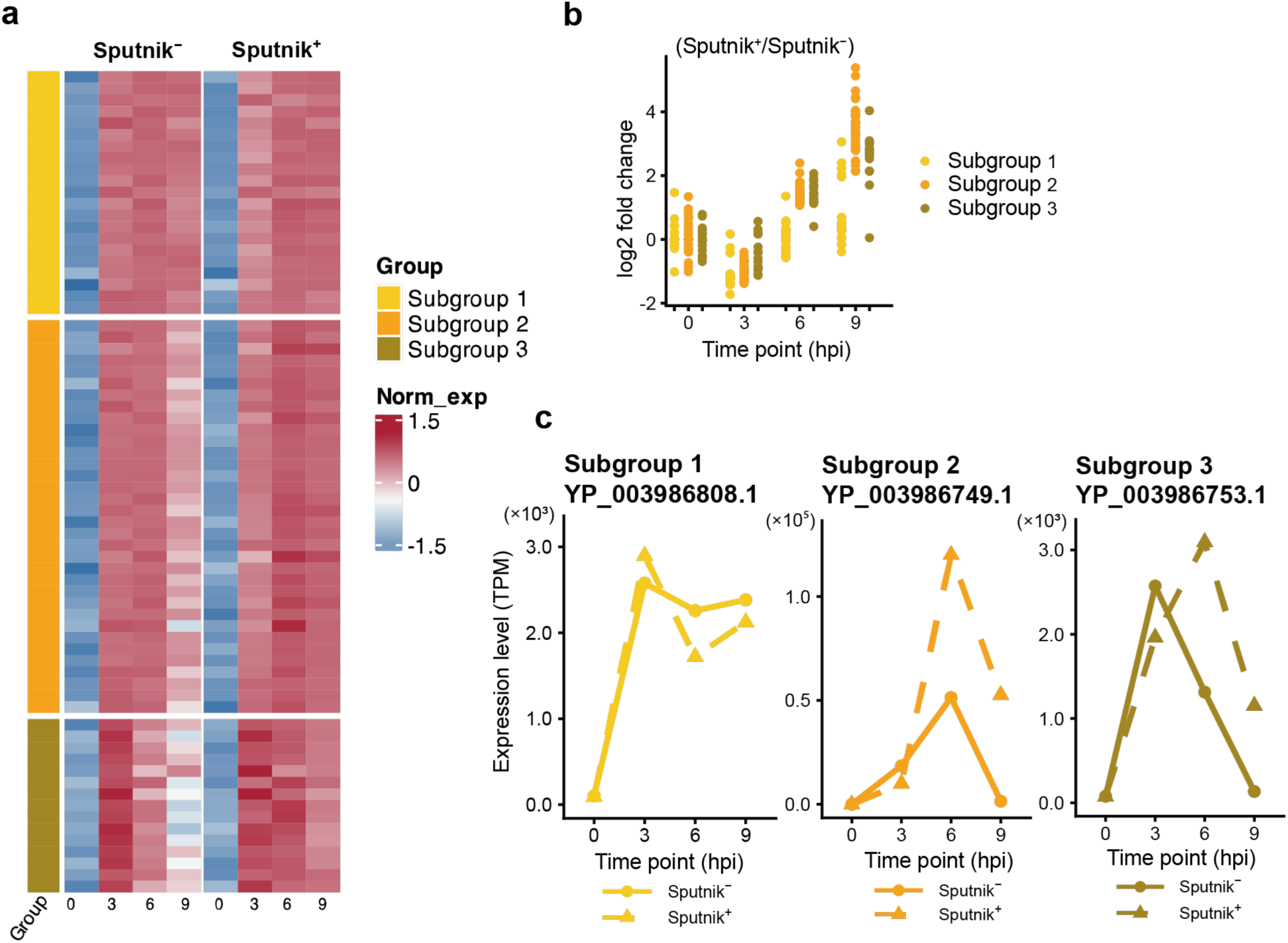
Expression profile of APMV intermediate genes. (a) Clustered heatmap showing normalized gene expression of APMV intermediate genes in Sputnik^−^ and Sputnik^+^ cells. Gene expression was calculated as TPM, then further normalized and clustered as described in Fig.6 (b) Expression changes from Sputnik^−^ to Sputnik^+^ cells at each time point. Each dot represents an APMV gene. (c) Temporal expression changes of representative genes. Genes with the highest expression from each subgroup are shown.

Late genes clustered into two subgroups (Fig. 9a). While both subgroups exhibited similar expression patterns, genes in subgroup 1 showed slightly higher expression levels than those in subgroup 2. At 0 hpi, there was no obvious difference between Sputnik^−^ and Sputnik^+^ cells. However, most genes showed lower expression in Sputnik^+^ cells compared to Sputnik^−^ cells at 3 and 6 hpi (Fig. 9b and c). This reduced expression was partially restored at 9 hpi, with some genes showing comparable expression levels in both conditions (Fig. 9b). These results suggest that Sputnik infection also delays the expression timing of APMV late genes.

**Fig 9.**
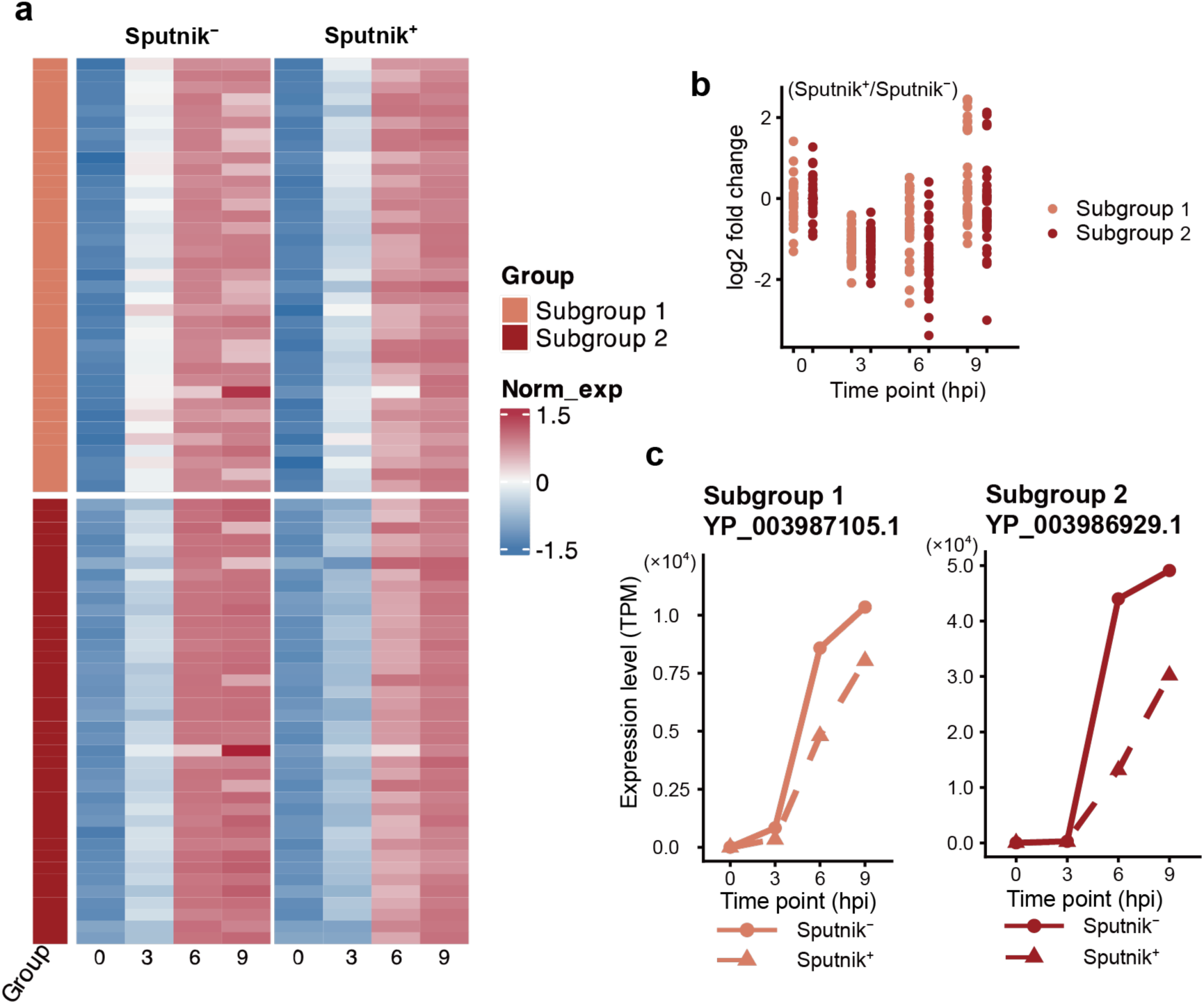
Expression profile for APMV late genes. (a) Clustered heatmap showing normalized gene expression of APMV late genes in Sputnik^−^ and Sputnik^+^ cells. Gene expressions was calculated as TPM, then further normalized and clustered as described in Fig.6 (b) Expression changes from Sputnik^−^ to Sputnik^+^ cells at each time point. Each dot represents an APMV gene. (c) Temporal expression changes of representative genes. Genes with the highest expression from each subgroup are shown.

### Promoter analysis of APMV and Sputnik genes

To investigate whether regulatory sequences contribute to the differential gene expression of APMV genes under Sputnik^+^ conditions, we searched for conserved motifs in the upstream regions of the APMV genes identified as differentially expressed. We identified only the previously reported conserved motif “AAAATTGA” in early expressed genes^19^ but failed to find specific promoter motifs (Supplementary Fig. 4). Additionally, a search for motifs in Sputnik genes revealed no statistically significant motifs (Supplementary Fig. 5).

## Discussion

Sputnik virophages infect mimiviruses and reduce their propagation^4^. However, the molecular mechanisms underlying this interference are largely unknown. In the present study, we investigated the transcriptome landscape of amoeba cells infected with APMV and Sputnik. We found that Sputnik infection drastically alters the APMV gene expression pattern at the late stages of infection. The PCA of the APMV gene expression profile indicated that Sputnik infection inhibits the progression of mimivirus infection from 6 to 9 hpi. The expression patterns of late and intermediate genes further support this delay. These genes exhibited reduced expression at the early stages of infection but reached similar or even higher expression levels at the late stages in Sputnik-infected cells compared to cells only infected with APMV. Taken together, our results demonstrate that Sputnik infection disrupts the transcriptional regulation of APMV at the late stages of infection.

Sputnik infection affects not only intermediate and late genes, but also immediate-early and early genes. The expression of these genes typically starts at the early stages of infection and generally decreases at the late stages. Although these genes also showed decreased expression at the late stages in Sputnik-infected cells, the decrease was smaller compared to cells without Sputnik infection. Consequently, higher expression levels were maintained at the late stages of infection in Sputnik-infected cells compared to those only infected with APMV. These results indicate that Sputnik infection prolongs the expression of APMV immediate-early and early genes. This prolongation suggests that Sputnik infection may hinder the transition from the early to the late stages of APMV infection, disrupting APMV late gene expression (Fig 10).

**Fig 10.**
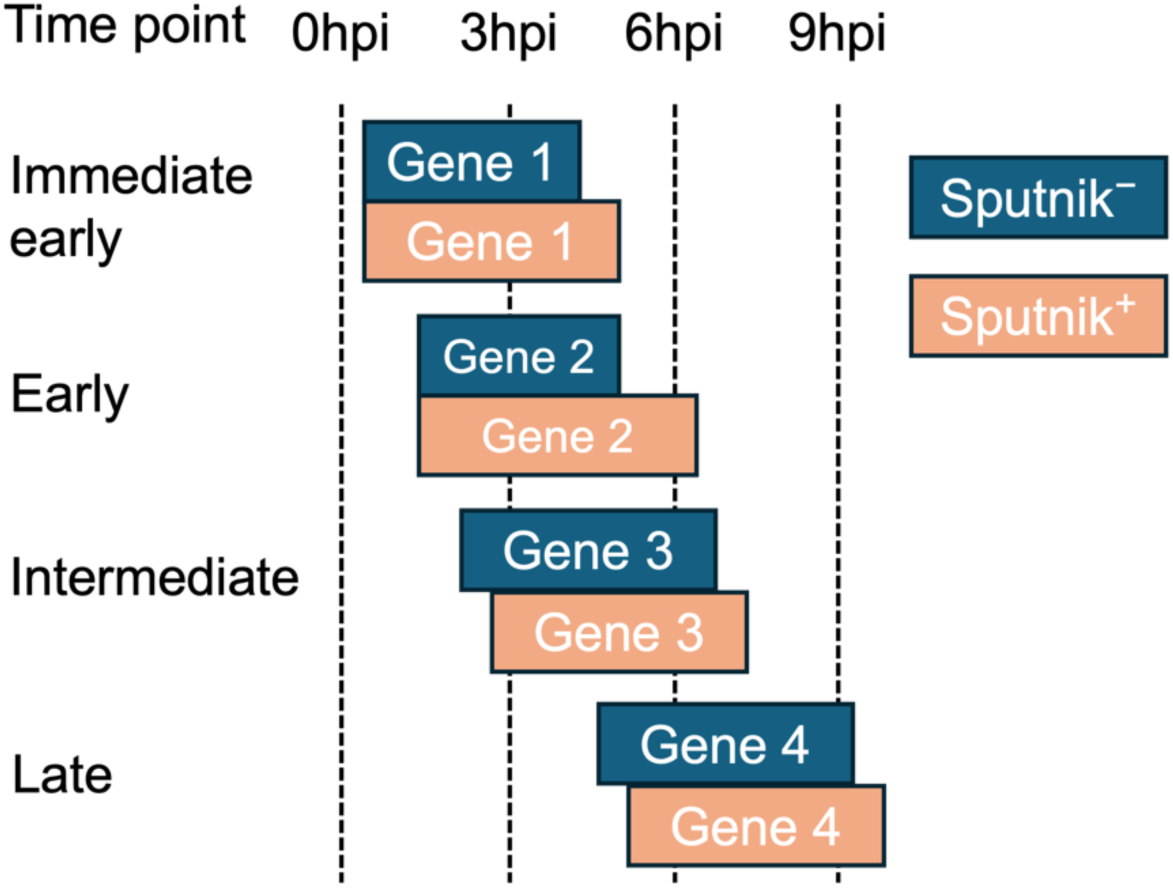
A hypothetical model of how Sputnik infection affects APMV gene expression Each box represents a period of APMV gene expression.

In Sputnik-infected cells, some genes exhibit expression levels similar to those in cells infected only with APMV, suggesting that Sputnik may target specific genes. APMV immediate-early and early genes typically include those involved in DNA replication. Prolonged expression of these genes might benefit Sputnik, as it possesses a limited number of genes related to DNA replication. However, since most APMV genes have not been fully characterized, further studies are needed to understand the advantage for Sputnik in disturbing APMV gene expression.

Since the discovery of virophages, it has been debated whether virophages are merely satellite viruses using mimiviruses as helper^20^ or if they are viruses that infect mimiviruses^21^. In this study, we revealed that Sputnik gene expression begins following APMV gene expression. Additionally, the sum of the viral-read proportion was similar between cells infected with and without Sputnik. These findings suggest that Sputnik exploits the transcription machinery of APMV. We also found that Sputnik infection has little effect on both the proportion of amoeba-derived reads and the amoeba gene expression pattern. Taken together, our results indicate that mimiviruses are primary hosts of Sputnik rather than helpers, and Sputnik uses amoeba as a helper.

The mechanisms by which Sputnik exploits APMV transcriptional machinery are currently unknown. We found that Sputnik does not necessarily share promoter motif with APMV. Additionally, expression changes caused by Sputnik varied even among genes expressed at the similar timing. These results imply that Sputnik can hijack specific transcriptional machinery to get precise manipulation of APMV genes. Combined with recent findings from other model systems of host giant virus-virophage interactions^17^, future studies may provide insights into the molecular mechanisms underlying this complex tripartite system.

## Materials and Methods

### Cells and viruses

*A. castellanii* (Douglas) Page, strain Neff (ATCC 30010), was maintained in peptone-yeast extract-glucose (PYG) medium at 28°C. APMV and Sputnik 3^4^ served as prototypes of mimivirus and virophage, respectively.

### Titration of viruses

The titer of APMV was measured using the 50% tissue culture infectious dose (TCID_50_) method on a 96-well plate. The titer of Sputnik was determined using a modified TCID_50_ method. Briefly, amoeba cells were infected with APMV at a multiplicity of infection (MOI) of 1 on a 96-well plate, followed by inoculation with a serially diluted Sputnik solution. Wells containing infectious Sputnik were initially inferred by observing a reduced cytopathic effect under light microscopy compared to wells containing only APMV. The presence of Sputnik was confirmed by quantitative polymerase chain reaction (qPCR) as follows: the entire volume of supernatant and remaining cells was collected from wells at the highest dilution level containing infectious Sputnik, as well as from one-order higher or lower dilution levels. Viral DNA was extracted from these wells using the following method: the collected supernatant was centrifuged at 15,000 rpm (Thermo Scientific, Sorvall ST 8FR centrifuge) for 1 hour at 4°C. The pellets were resuspended in 45 µl of 50 mM sodium hydroxide solution and incubated at 95°C for 5 minutes. Then, 5 µl of 1 M Tris(hydroxymethyl)aminomethane hydrochloride (pH 8.0) and 450 µl Tris-EDTA buffer (10 mM Tris, 1 mM EDTA, pH8.0) were added to the solution, which was then subjected to qPCR. The reaction mixture was prepared using the KAPA PROBE Fast qPCR kit (ROXLowqPCR, KAPA BIOSYTEMS), with 0.3 µM forward and reverse primers and 0.4 µM probe. The PCR conditions were as follows: 95°C for 3 min; 40 cycles of 95°C for 3 sec, 60°C for 20 sec, and 72°C for 1 sec. The sequences of the primers and probe^22^ are shown in Supplementary Table 4.

### Sample preparation

A total of 1 × 10^6^ amoeba cells were inoculated with APMV or APMV and Sputnik at an MOI of 10 in PYG medium. The inoculated cells were incubated for 1 h at room temperature with shaking. After incubation, the culture medium was replaced with 1 ml of phosphate buffered saline (PBS) to remove uninfected viruses. The PBS was then replaced with 1.5 ml of fresh PYG medium, and the cells were incubated at 30°C for the designated periods. The time point at which the medium was exchanged for fresh PYG medium was designated as 0 hpi.

### RNA extraction

Infected cells were collected at 0, 1, 3, 6, 9, and 12 hpi using a cell scraper (Thermo Fisher Scientific) and then centrifuged at 2,000 rpm for 5 min at 4°C. The cell pellet was stored in 1 ml of TRIzol (Invitrogen) at −80°C. Three independent experiments were conducted as biological replicates. RNA extraction followed the manufacture’s protocol. Briefly, 20% of the sample volume of chloroform was added to the cells in TRIzol. The RNA-containing aqueous layer was transferred to new tubes, to which 500 ml of isopropanol was added to precipitate RNA. The RNA pellet was washed with 70% ethanol, dissolved in nuclease-free water and treated with DNase I (New England Biolabs). RNA was then re-extracted using phenol/chloroform and re-precipitated with 99.5% ethanol and 0.3 M sodium acetate. The precipitate was washed with 70% ethanol and dissolved in nuclease-free water. RNA concentration was measured using a Qubit 4 Fluorometer (Invitrogen) with the Qubit^TM^ RNA BR Assay Kit (Invitrogen). RNA purity was assessed by measuring the A260/A280 and A260/A230 ratio using a DeNovix DS-11 spectrophotometer (SCRUM Inc.).

### RNA sequencing

Quality control and sequencing of RNA samples were performed by Rhelixa Inc. (Japan). In brief, strand-specific libraries were prepared using poly-A selection and sequenced to a depth of 5 G bases per sample using Illumina NovaSeq 6000.

### Measuring gene expression by qPCR

Extracted RNA was adjusted to a uniform concentration across all samples using nuclease-free water. For cDNA synthesis, 0.5 µg of total RNA was used with D(T)23VN primer (New England Biolabs), 10 mM dNTPs (New England Biolabs), AMV reverse transcriptase (New England Biolabs), and a RNase inhibitor (New England Biolabs). The reaction was conducted at 42°C for 1h, followed by enzyme deactivation at 80°C for 5 min.

The synthesized cDNA was then subjected to qPCR using the KAPA SYBR Fast qPCR kit (ROXLowqPCR, KAPA BIOSYTEMS) with 0.2 µM of forward and reverse primers. The sequences of primers^22,23^ are listed in Supplementary Table 4. The PCR conditions were as follows: 95°C for 20 sec; 40 cycles of 95°C for 15 sec, 60°C for 20 sec and 72°C for 10 sec. Relative gene expression was calculated using the 2^ΔCt^ method, with the highest expression level set as 100.

### Data analysis

The quality of reads was assessed using FastQC (v0.12.0)^24^. With an overall quality score exceeding 20 and no known adaptors detected, no trimming was performed. Reads were mapped to a sequence dataset consisting of the Sputnik 3 genome (NC_011132.1) and the APMV genome (NC_014649.1) using HISAT2 (v 2.2.1), ^25^ with a maximum intron size of 5,000 bp. Unmapped reads were subsequently mapped to the *A. castellanii* genome (GCF_000313135.1_Acastellanii.strNEFF_v1) with a maximum intron size of 500,000 bp. Output data were processed with Samtools (v1.19)^26^. The number of reads mapped to each gene was counted using HTSeq (v2.0.5)^27^ in union mode with the reverse strand-specific assay option. Gene expression levels were normalized to transcripts per million (TPM).

PCA and DEG detection were conducted using DESeq2 (1.42.0)^28^. Genes were classified as DEGs if they had a False Discovery Rate (FDR) < 0.05 and absolute log2-fold change ≥ 1^29^.

Clustering was performed for genes meeting two criteria: FDR < 0.05 at every time point and an absolute log2-fold change ≥ 1 at least one time point. Selected genes were clustered using the k-means method based on TPM values normalized by adjusting the mean to 0 and the variance to 1 for each gene. The optimal number of clusters for Sputnik and APMV was determined using the total-within-sum-of-squares method in factoextra (v1.0.7).

Annotation information for APMV genes was retrieved from the National Center for Biotechnology Information GenBank database. Functional categories for each APMV gene were assigned manually, referring to previous studies^19,30–32^.

### Promoter search

One hundred bp sequences upstream of the open reading frames of each DEG in Fig 5 matching the same expression timing were extracted. Sequence motifs were predicted using MEME 5.5.4^33^. MEME was run in classic mode with motif width ranges set from 8 to 25 bp and the “Any Number of Repetitions” option.

### Visualization

All data processing was carried out using the dplyr (v1.0.10)^34^ and tidyr (v1.3.0)^35^ packages in R (v4.2.1)^36^. Heatmaps were generated with the pheatmap (v1.0.12) package^37^. All plots were created using the ggplot2 (v3.4.4) package^38^.

## Data availability

The RNA sequencing data are available at DDBJ under accession number PRJDB18584.

## Acknowledgements

We thank Prof. Bernard La Scola and Ms. Lina Barrassi for kindly providing APMV and Sputnik 3 and Dr. Matthias Fischer and Dr. Anna Koslová for kindly sharing the method of virophage titration. Computation time was provided by the SuperComputer System, Institute for Chemical Research, Kyoto University. This study was supported by the Japan Society for the Promotion of Science KAKENHI grant numbers 22K15175 to HH and 22H00384 to OH and by Japan Science and Technology Agency ACT-X Grant number JPMJAX22BI to HH.

## Author Contributions

JC, HO, and HH conceptualized the study. JC performed experiments and formal analysis. JC and HH wrote the manuscript with intellectual input from all authors. HH and OH supervised the project.

## Reference

1. Duponchel, S. & Fischer, M. G. Viva lavidaviruses! Five features of virophages that parasitize giant DNA viruses. PLOS Pathog. 15, e1007592 (2019).

2. Dance, A. Beyond coronavirus: the virus discoveries transforming biology. Nature 595, 22– 25 (2021).

3. Koonin, E. V., Senkevich, T. G. & Dolja, V. V. The ancient Virus World and evolution of cells. Biol. Direct 1, 29 (2006).

4. La Scola, B. et al. The virophage as a unique parasite of the giant mimivirus. Nature 455, 100–104 (2008).

5. Claverie, J.-M. & Abergel, C. Mimivirus and its virophage. Annu. Rev. Genet. 43, 49–66 (2009).

6. Fischer, M. G. & Suttle, C. A. A virophage at the origin of large DNA transposons. Science 332, 231–234 (2011).

7. Roitman, S. et al. Isolation and infection cycle of a polinton-like virus virophage in an abundant marine alga. Nat. Microbiol. 2023 82 8, 332–346 (2023).

8. Paez-Espino, D. et al. Diversity, evolution, and classification of virophages uncovered through global metagenomics. Microbiome 7, 157 (2019).

9. Coutinho, F. H., Gregoracci, G. B., Walter, J. M., Thompson, C. C. & Thompson, F. L. Metagenomics Sheds Light on the Ecology of Marine Microbes and Their Viruses. Trends Microbiol. 26, 955–965 (2018).

10. Zhou Jinglie et al. Diversity of Virophages in Metagenomic Data Sets. J. Virol. 87, 4225– 4236 (2013).

11. Oh, S., Yoo, D. & Liu, W.-T. Metagenomics Reveals a Novel Virophage Population in a Tibetan Mountain Lake. Microbes Environ. 31, 173–177 (2016).

12. Yutin, N., Kapitonov, V. V. & Koonin, E. V. A new family of hybrid virophages from an animal gut metagenome. Biol. Direct 10, 19 (2015).

13. Tokarz-Deptuła, B., Czupryńska, P., Poniewierska-Baran, A. & Deptuła, W. Characteristics of virophages and giant viruses. Acta Biochim. Pol. (2018) doi:10.18388/abp.2018_2631.

14. Mougari, S. et al. A virophage cross-species infection through mutant selection represses giant virus propagation, promoting host cell survival. *Commun*. Biol. 3, 248 (2020).

15. Mougari, S. et al. Guarani Virophage, a New Sputnik-Like Isolate From a Brazilian Lake. Front. Microbiol. 10, 1003 (2019).

16. Gaia, M. et al. Zamilon, a Novel Virophage with Mimiviridae Host Specificity. PLoS ONE 9, e94923 (2014).

17. Fischer, M. G. & Hackl, T. Host genome integration and giant virus-induced reactivation of the virophage mavirus. Nature 540, 288–291 (2016).

18. Boyer, M. et al. Mimivirus shows dramatic genome reduction after intraamoebal culture. Proc. Natl. Acad. Sci. 108, 10296–10301 (2011).

19. Legendre, M. et al. mRNA deep sequencing reveals 75 new genes and a complex transcriptional landscape in Mimivirus. Genome Res. 20, 664–674 (2010).

20. Hu, C.-C., Hsu, Y.-H. & Lin, N.-S. Satellite RNAs and Satellite Viruses of Plants. Viruses 1, 1325–1350 (2009).

21. Krupovic, M. & Cvirkaite-Krupovic, V. Virophages or satellite viruses? Nat. Rev. Microbiol. 9, 762–763 (2011).

22. Ngounga, T. et al. Real-Time PCR Systems Targeting Giant Viruses of Amoebae and Their Virophages. Intervirology 56, 413–423 (2013).

23. Gaia, M., et al. Broad Spectrum of Mimiviridae Virophage Allows Its Isolation Using a Mimivirus Reporter. PLoS ONE 8, e61912 (2013).

24. Andrews, S. FastQC: A Quality Control Tool for High Throughput Sequence Data [Online]. (2010).

25. Kim, D., Paggi, J. M., Park, C., Bennett, C. & Salzberg, S. L. Graph-based genome alignment and genotyping with HISAT2 and HISAT-genotype. Nat. Biotechnol. 37, 907–915 (2019).

26. Danecek, P. et al. Twelve years of SAMtools and BCFtools. GigaScience 10, (2021).

27. Putri, G. H., Anders, S., Pyl, P. T., Pimanda, J. E. & Zanini, F. Analysing high-throughput sequencing data in Python with HTSeq 2.0. Bioinforma. Oxf. Engl. 38, 2943–2945 (2022).

28. Love, M. I., Huber, W. & Anders, S. Moderated estimation of fold change and dispersion for RNA-seq data with DESeq2. Genome Biol. 15, 550 (2014).

29. Rodrigues, R. A. L. et al. Analysis of a Marseillevirus Transcriptome Reveals Temporal Gene Expression Profile and Host Transcriptional Shift. Front. Microbiol. 11, 651 (2020).

30. Tatusov, R. L. et al. The COG database: an updated version includes eukaryotes. BMC Bioinformatics 4, 41 (2003).

31. Koonin, E. V. & Yutin, N. Origin and Evolution of Eukaryotic Large Nucleo-Cytoplasmic DNA Viruses. Intervirology 53, 284–292 (2010).

32. Yutin, N., Wolf, Y. I., Raoult, D. & Koonin, E. V. Eukaryotic large nucleo-cytoplasmic DNA viruses: Clusters of orthologous genes and reconstruction of viral genome evolution. Virol. J. 6, 223 (2009).

33. Bailey, T. L., Johnson, J., Grant, C. E. & Noble, W. S. The MEME Suite. Nucleic Acids Res. 43, W39–W49 (2015).

34. Wickham H, François R, Henry L, Müller K, Vaughan D (2023). _dplyr: A Grammar of Data Manipulation_. R package version 1.1.4, <https://CRAN.R-project.org/package=dplyr>.

35. Wickham H, Vaughan D, Girlich M (2024). _tidyr: Tidy Messy Data_. R package version 1.3.1, <https://CRAN.R-project.org/package=tidyr>.

36. R Core Team. R: A Language and Environment for Statistical Computing. (R Foundation for Statistical Computing, Vienna, Austria, 2024).

37. Kolde R (2019). _pheatmap: Pretty Heatmaps_. R package version 1.0.12, <https://CRAN.R-project.org/package=pheatmap>.

38. Wickham, H. Ggplot2: Elegant Graphics for Data Analysis. (Springer-Verlag New York, 2016).

