## Supplementary Fig. 1, Supplementary Fig. 2, Supplementary Fig. 3, Supplementary Fig. 4, Supplementary Fig. 5, Supplementary Table 4 for "Sputnik virophage disrupts the transcriptional regulation of its host giant virus"

**Supplementary Fig. 1–5**

*** Supplementary Table 1. Counts of reads mapped to the Sputnik genome**

*** Supplementary Table 2. Differentially expressed host genes**

*** Supplementary Table 3. APMV genes with assigned clusters and annotations**

**Supplementary Table 4. List of primers**

**References**

(*Included in a separated Excel file.)

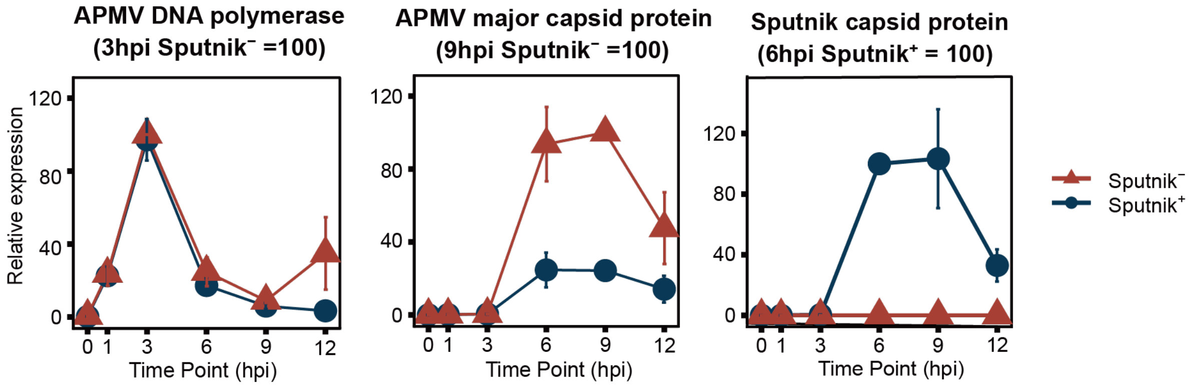


**Supplementary Fig 1. Expression profiles of representative APMV and Sputnik genes**.

Relative expression of representative genes was measured by quantitative polymerase chain reaction. Error bars represent the standard error (n = 3). Expression levels are normalized as follows: 100 at 3 hpi for APMV DNA polymerase in Sputnik$^{-}$ cells, 100 at 9 hpi for APMV major capsid protein in Sputnik$^{-}$ cells, and 100 at 6 hpi for Sputnik capsid protein in Sputnik$^{+}$ cells.

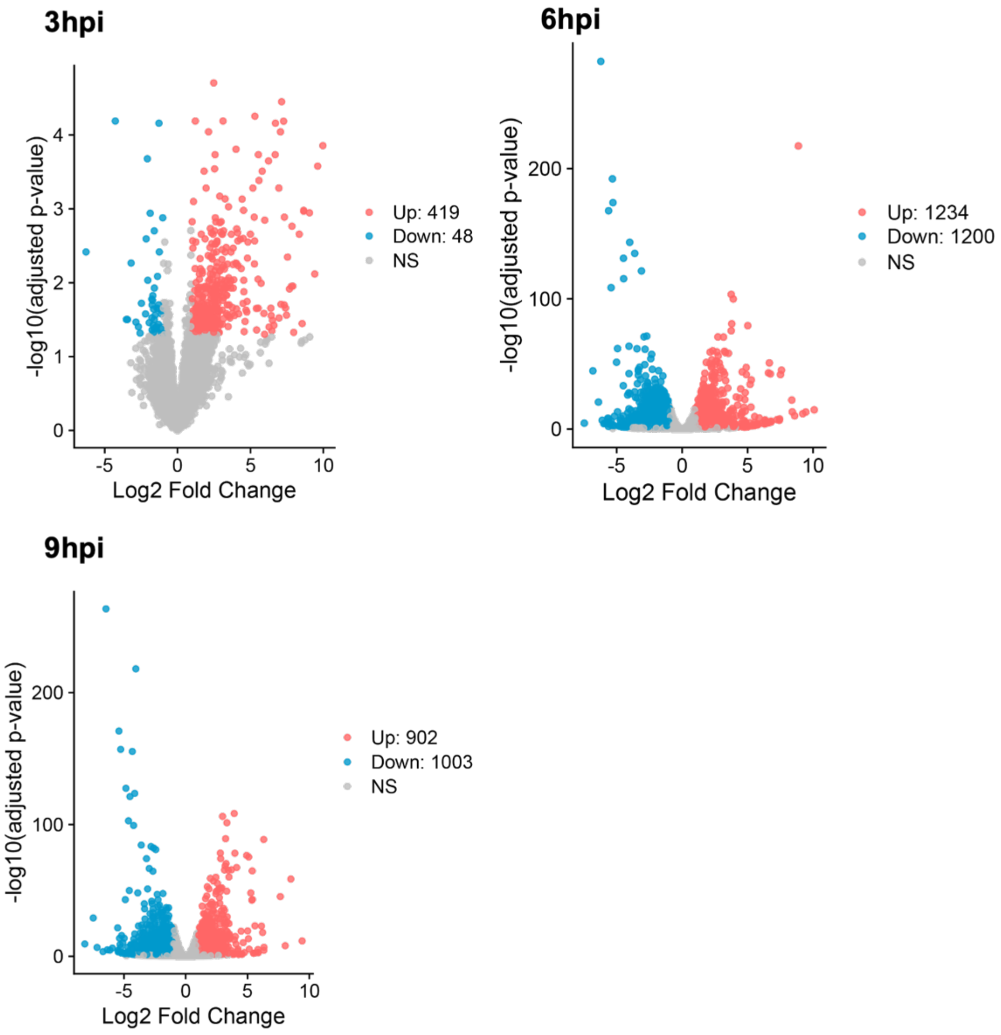


**Supplementary Fig 2. Comparison of amoeba gene expression in Sputnik⁻ cells over time.**

Gene expression in Sputnik⁻ cells at 3, 6, and 9 hpi was compared to expression at 0 hpi. Genes with $p_{adj} \leq 0.05$ and $|\log2 \text{ fold change}| \geq 1$ are considered DEGs and are marked in red or blue. Red and blue represent up-regulated and down-regulated genes, respectively. NS denotes no significant difference.

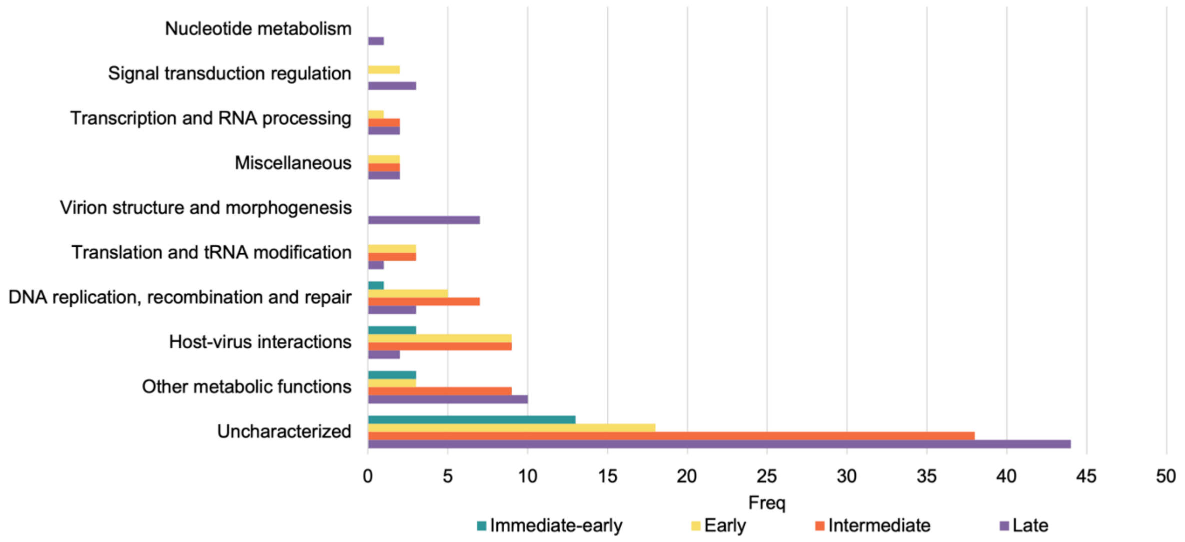


**Supplementary Fig 3. Functional annotations of APMV genes across different clusters.**

The number of genes assigned to each functional category is displayed. Each color indicates the timing of gene expression. Functional categories for each APMV gene were manually assigned based on previous studies[1–5].

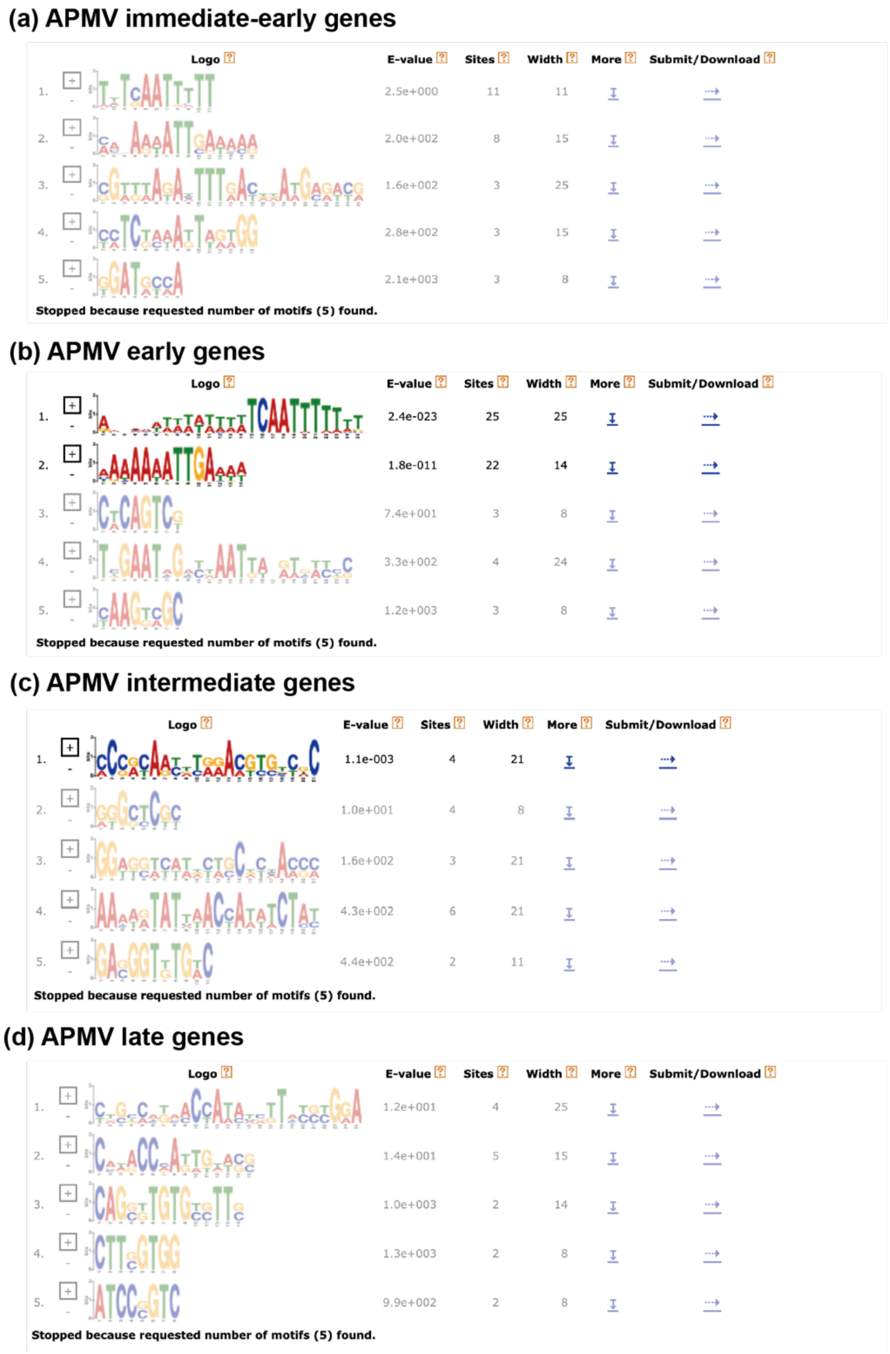


**Supplementary Fig 4. Promoter analysis of APMV using MEME.**

Predicted motifs in (a) immediate-early, (b) early, (c) intermediate, and (d) late genes with differential expression (false discovery rate < 0.05 and |log2-fold change| ≥ 1). Motifs that met the significance threshold (E-value < 0.05) highlighted in bright colors.

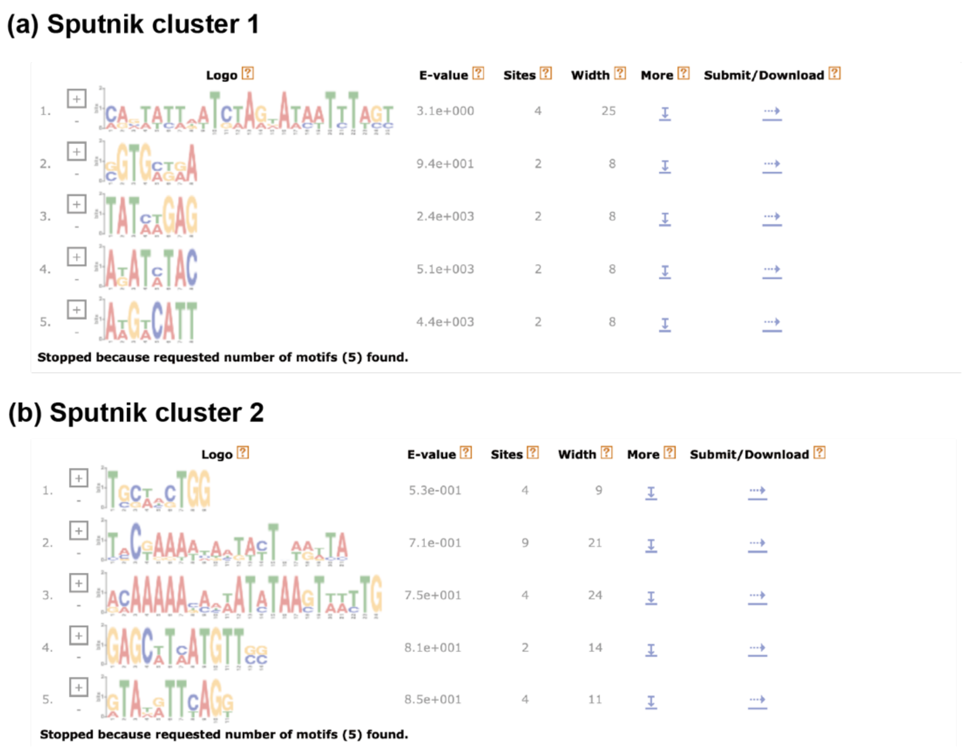


**Supplementary Fig 5. Promoter analysis of Sputnik using MEME.**

Predicted motifs for (a) early and (b) late Sputnik genes. No motifs met the significance threshold (E-value < 0.05).

### Supplementary Table 4. List of primers.

| Target | Sense | Sequence (5' → 3') |
|---|---|---|
| Sputnik capsid protein | Forward | GAGATGCTGATGGAGCCAAT |
| | Reverse | CATCCCACAAGAAAGGAGGA |
| APMV DNA polymerase | Forward | TGCGGGAGTTGGAGAAATGATTGTC |
| | Reverse | TTGGCAGCCCTTTGACACTTC |
| APMV major capsid protein | Forward | GAACCTGGAGGTTATGAATGTGAAGG |
| | Reverse | ACCATCGAAAGCTTCAGCAGTGG |
| Sputnik probe | | TACTTCAGCAGCTGGTCTTTCTGA |

### References


[1] Boyer, M. *et al.* Mimivirus shows dramatic genome reduction after intraamoebal culture. *Proc. Natl. Acad. Sci.* 108, 10296–10301 (2011).

[2] Legendre, M. *et al.* mRNA deep sequencing reveals 75 new genes and a complex transcriptional landscape in Mimivirus. *Genome Res.* 20, 664–674 (2010).

[3] Tatusov, R. L. *et al.* The COG database: an updated version includes eukaryotes. *BMC Bioinformatics* **4**, 41 (2003).

[4] Koonin, E. V. & Yutin, N. Origin and Evolution of Eukaryotic Large Nucleo-Cytoplasmic DNA Viruses. *Intervirology* **53**, 284–292 (2010).

[5] Yutin, N., Wolf, Y. I., Raoult, D. & Koonin, E. V. Eukaryotic large nucleo-cytoplasmic DNA viruses: Clusters of orthologous genes and reconstruction of viral genome evolution. *Virol. J.* **6**, 223 (2009).
